# Unbiased Quantification of Persistent Postural and Motor Deficits Following Spinal Cord Injury in Mice

**DOI:** 10.1101/2025.05.24.655880

**Authors:** Sarah S. Jobbins, Vihaan Pande, Chih-Wei Zeng, Helen Poldsam, Allan-Hermann Pool, Chun-Li Zhang, Helen C. Lai

## Abstract

Spinal cord injury (SCI) causes multifaceted postural and motor impairments that are challenging to quantify. Conventional behavioral tests, such as the Basso mouse scale (BMS), rely on qualitative observations, which do not detect subtle yet significant functional deficits. To perform unbiased and quantitative evaluation of motor function and posture after SCI, we employed high-speed video tracking with machine learning-based whole body pose estimation and weight-bearing analyses using the Blackbox system in freely moving mice. We found enduring alterations in posture induced by SCI not captured by conventional metrics. Postural deficits included reduced hindpaw spacing with concomitant increased forepaw spacing, narrowed femur width, and altered hind paw angles, which remained evident beyond 42 days post-injury (dpi). Furthermore, sustained deficits in locomotor activity were identified as decreased distance traveled, decreased exploratory behavior, and disrupted fore-to-hindpaw speed ratio. Additionally, we analyzed behavioral motifs using Keypoint MoSeq software and found frequency changes in unique motor syllables correlating with forward acceleration and turning after SCI. Interestingly, while motor deficits persisted, sensory deficits, such as thermal and mechanical sensitivity, returned to baseline levels by 21 days after injury in C57Bl/6J mice, with no subsequent hypersensitivity. Lastly, we developed a web-based application to assist in visualization and analyses of Blackbox-based kinematic data. Altogether, our study identifies distinct postural and motor deficits pre- and post-SCI using accurate, unbiased, and quantitative behavioral assessments. By tracking the unique features of motor recovery trajectories, researchers can more accurately assess the effectiveness of SCI therapeutic strategies.

## Introduction

In humans, SCI typically results in persistent impairments in both motor and sensory function (Awai & Curt, 2014; Finnerup et al., 2001; Siddall & Loeser, 2001). Mice are a widely used model system for spinal cord injury (SCI) due to the ease of testing pharmacological agents and genetically tractable interventions. A unique feature of SCI in mouse models is their apparent spontaneous recovery of locomotor function over time, which can obscure subtle but functionally significant deficits in coordination, weight-bearing, and fine motor control. Traditional methods for assessing motor recovery in mice often rely on subjective observations and may fail to detect these nuanced impairments. This lack of precision limits our ability to fully characterize the extent of recovery or to quantitatively assess incomplete recovery, which is critical for evaluating the effectiveness of therapeutic interventions (Fawcett et al., 2007).

Motor behavioral assays commonly used in SCI research include Basso mouse scale (BMS) locomotor scale, open field gait analysis, the ladder rung test, and the CatWalk automated gait analysis. Each of these tools has specific limitations. The BMS scale relies on subjective scoring, which introduces inter-rater variability and reduces reproducibility (Basso et al., 2006). Open field tests primarily assess gross locomotor activity and anxiety-related behaviors, such as total distance traveled, velocity, and time spent in the center versus periphery, but provide limited resolution for evaluating coordination, postural control, or limb-specific deficits in gait (Gould et al., 2009; Walsh & Cummins, 1976). The ladder rung and CatWalk tests capture paw placement accuracy and gait parameters, respectively, along a predefined, constrained pathway, limiting their ability to assess natural gait dynamics and subtle impairments in posture or coordination (Cummings et al., 2007; Gabriel et al., 2009; Hamers et al., 2006; Metz & Whishaw, 2002).

These challenges underscore the need for more unbiased and multidimensional methods for assessing motor recovery in SCI models. To address these limitations, we implemented an advanced kinematic and weight-bearing analysis platform (Blackbox One) that provides a feature-rich and objective evaluation of motor function in freely moving mice. This system integrates high-speed video tracking with machine learning-based body part tracking and whole body pose estimation and weight-sensing based on frustrated total internal reflection (FTIR) optical technology to analyze detailed body positioning and weight distribution of a freely moving mouse (Zhang et al., 2022). By quantifying key movement parameters such as joint angles, paw luminance ratios, and skeletal symmetry, the Blackbox system enables the detection of stereotypic behavioral features, subtle motor impairments and incomplete recovery trajectories that are often overlooked by conventional techniques. Furthermore, its automated data collection and processing minimize variability associated with subjective behavioral scoring, enhancing reproducibility across experiments (Zhang et al., 2022).

In this study, we show that automated kinematic tracking and weight-bearing analysis can detect previously undetected alterations in postural adaptation, motor coordination, and weight distribution during SCI recovery. In addition, we found that the postural adaptations detected with the Blackbox system were not due to sensory hypersensitivity. By leveraging this analytical platform, we aim to improve the precision of SCI motor measurements, which will facilitate the evaluation of novel therapeutic strategies and bridge the gap in motor assessments between preclinical models and clinical applications.

This work highlights the importance of integrating high-resolution motor analysis into SCI research, ultimately contributing to the development of more effective treatment strategies for individuals with spinal cord injuries.

## Results

### Characterization of the SCI Model and Injury Lesion

In a T9 contusion SCI model, we assessed motor and sensory recovery in female C57Bl/6J mice one day post-injury (1 dpi) and then every week up to 6 weeks post-SCI compared to sham controls (Figure 1A). To validate our contusion SCI model, we conducted histological analyses at 42 dpi using immunohistochemistry (IHC) (Figure 1B-D). We determined the extent of tissue damage in transverse spinal cord sections stained with DAPI (nuclear marker), GFAP (astrocyte marker), and PDGFRβ (fibroblast marker). GFAP staining marks the glial scar, which plays roles in both neuroprotection and regeneration inhibition (Silver & Miller, 2004), and PDGFRβ staining marks the fibrotic scar, which forms a dense extracellular matrix barrier that inhibits axonal regrowth (Anderson et al., 2016). Representative images from sham and SCI groups highlight substantial structural differences (Figure 1B). The sham control group exhibited well-preserved spinal cord architecture with uniform GFAP expression, whereas the SCI group demonstrated a prominent fibrotic lesion core with a significant accumulation of PDGFRβ+ fibroblasts encased by a dense GFAP+ astroglial scar. Quantitative analysis of the injury area in four animals, revealed a significantly larger fibrotic lesion in the SCI group compared to sham controls (*p*< 0.0001) (Figure 1C), confirming extensive tissue damage post-injury.

**Figure 1.**
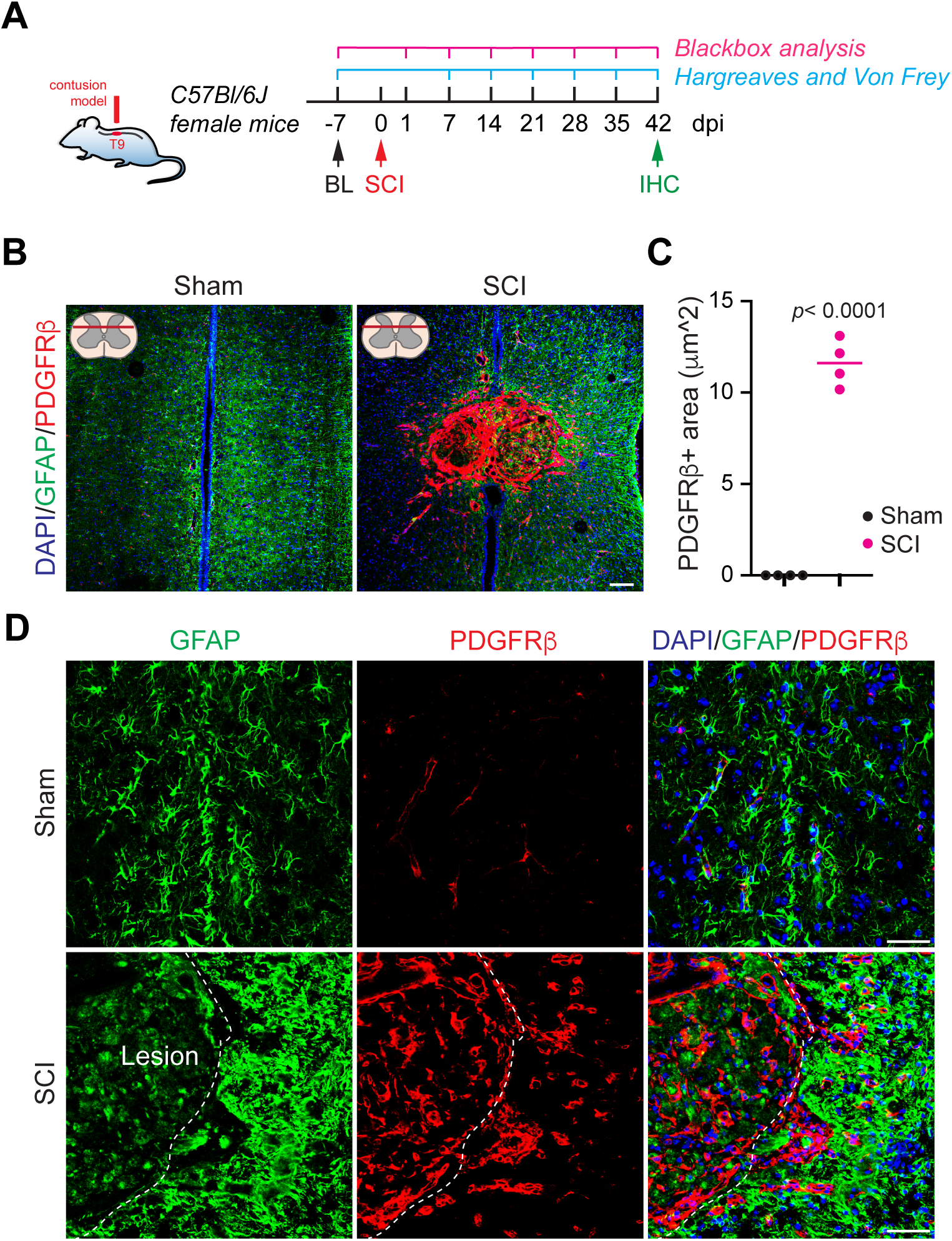
SCI induces astrocyte and fibroblast activation, leading to glial and fibrotic scar formation. (A) Experimental schematic: Wild-type mice were subjected to SCI at day 0, and spinal cord tissue was analyzed at 42 dpi using IHC. Hargreaves and von Frey behavioral assays were assessed at baseline (BL) (7 days prior to SCI) and every 7 days for 6 weeks. Blackbox was assessed at BL, 1 dpi and 7 day intervals for 6 weeks. (B) Representative transverse spinal cord sections stained with GFAP (green, astrocytes), PDGFRβ (red, fibroblasts), and DAPI (blue, nuclei). Sham-operated mice exhibit preserved spinal cord structure, while SCI sections demonstrate a prominent lesion core marked by increased GFAP and PDGFRβ expression. Scale bar = 100 µm. (C) Quantitative analysis of the PDGFRβ+ injury area (µm²) indicates a significantly larger lesion size in the SCI group compared to Sham controls (\*\*\*\**p*< 0.0001, two-tailed unpaired *t*-test, n=4 per group). Data corresponds to the largest PDGFRβ area such as the representative SCI image in (B). (D) Higher magnification views of the lesion microenvironment. In SCI conditions, GFAP-positive astrocytes (green) form a glial scar encasing the lesion site, and PDGFRβ-positive fibroblasts (red) are enriched within the lesion core, contributing to fibrotic scar formation. The dashed line outlines the lesion boundary. DAPI (blue) highlights the distribution of nuclei. Scale bar = 50 µm.

Further investigation into the lesion microenvironment (Figure 1D) revealed pronounced astroglia reactivity, as evidenced by elevated GFAP expression outlining the lesion site, and a high enrichment of PDGFRβ+ fibroblasts within the lesion core. The distinct boundary between GFAP+ reactive astrocytes forming the glial scar and PDGFRβ+ fibroblasts contributing to the fibrotic scar underscores the dual nature of the injury response and aligns with established SCI pathology (Ayazi et al., 2022; Tran et al., 2022). These findings confirm the successful establishment of a contusion SCI model, characterized by hallmark histopathological features of fibrotic and glial scar formation.

### Mice Exhibit Postural Adjustments during Motor Recovery

Postural impairments following SCI were revealed using machine learning-based whole body pose estimation tracking data with the Blackbox system (Figure 2A). Analysis of the skeleton-based tracking features revealed several parameters in posture and locomotion that were significantly different upon SCI (Figure 2B). These include distance between various body parts, paw and limb angles, and distance traveled. We followed up on several of these features to understand how posture and locomotion change post-SCI.

**Figure 2.**
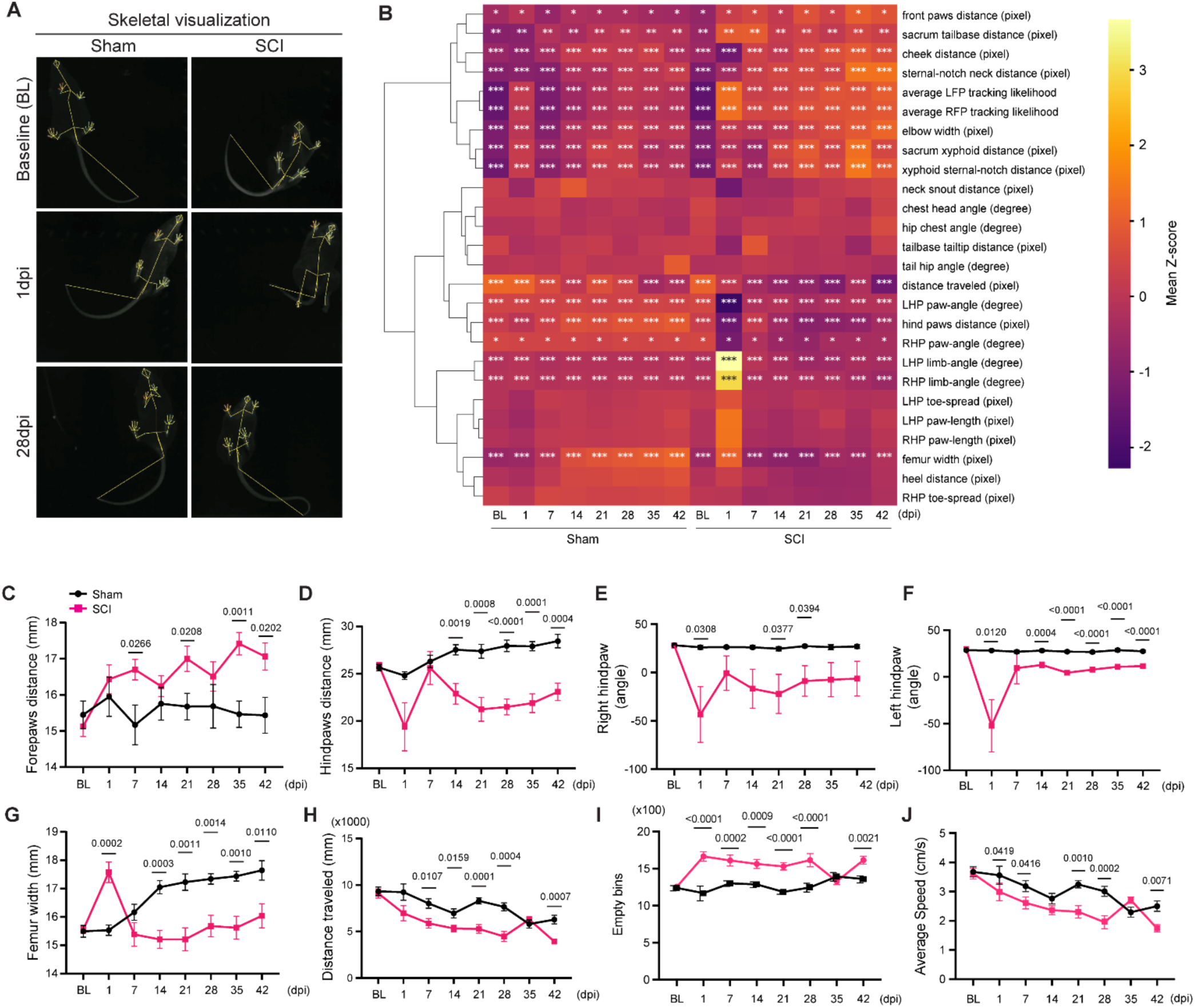
Changes in animal posture and locomotion following SCI. (A) Representative skeletal visualizations of animal posture at baseline (BL), 1-day post-injury (dpi), and 28 dpi in Sham and SCI-treated animals. Skeletal representations illustrate limb positioning and coordination changes over time. (B) Hierarchical heatmap of the mean z-score for all detected features utilizing pose estimation skeleton tracking. Significance is represented as *** = *p*<0.001, ** = *p*<0.01, and * = *p*<0.05. A sample of the significantly different postural and motor parameters are: (C) Distance between forepaws; (D) Distance between hindpaws; (E, F) Right and Left Hindpaw angles, measured as the angle between the intersection of a vector from the tail base to the sacrum and a vector from the heel to the center of the paw; (G) Femur width, (H) Total distance traveled during the recording session, representing overall locomotor activity. Tracking data was also used to analyze additional parameters: (I) Empty bin counts reflecting exploratory behavior and movement patterns; (J) Average speed in SCI animals over time. A two-way ANOVA is used in (B). A two-tailed unpaired Student’s t-test for comparisons between Sham and SCI at specific timepoints is used in (C-J). Abbreviations: Left and Right Forepaw (LFP, RFP), Left and Right Hindpaw (LHP, RHP).

At baseline, both sham and SCI animals displayed similar postural alignment. However, by 1 dpi, SCI-treated animals exhibited noticeable asymmetry. SCI-treated animals consistently positioned their forepaws farther apart compared to sham animals, with a significant increase observed at 7 dpi (*p*= 0.0266) that persisted through 42 dpi (*p*= 0.0202; Figure 2C), likely reflecting a compensatory mechanism to shift weight forward and stabilize the forelimbs due to hindlimb deficits. Conversely, the distance between hindpaws significantly decreased at 14 dpi (*p*= 0.0019) and remained diminished compared to sham controls throughout the 42-day study (*p*= 0.0004; Figure 2D). Similarly, femur spacing initially increased at 1 dpi (*p*= 0.0002) but subsequently decreased by 14 dpi (*p*= 0.0003), with reductions persisting through 42 dpi (*p*= 0.0110; Figure 2G), indicating restricted hindlimb range of motion. Additionally, both left and right hindpaw angles (hindpaw-tail base-sacrum angle) were significantly reduced in SCI-treated animals relative to sham controls. The right hindpaw angle showed a significant reduction at 1 dpi (*p*= 0.0308) and exhibited partial recovery over time, though transient reductions remained significant throughout recovery (Figure 2E). Similarly, the left hindpaw angle was significantly reduced at 1 dpi (*p* = 0.0120) and, despite partial recovery, remained significantly lower at 42 dpi (*p*< 0.0001; Figure 2F). These findings suggest a compensatory lateral rotation of the hindpaws post-SCI.

We note that we performed this analysis with the original version of the Blackbox machine learning-based whole body post estimation software. With this version, we observed that the software made transient errors in pose estimation during the acute phase of SCI recovery (Supplemental Figure 1). Blackbox continues to improve the DeepLabCut model and future versions of the software will take SCI models into account.

### Changes in Locomotor Activity and Coordination during Motor Recovery

Body part tracking data also revealed changes in locomotor activity and coordination. The total distance traveled in 5 min. was substantially decreased at 7 dpi compared to sham controls (*p*= 0.0107) and remained until 42 dpi (*p*= 0.0007; Figure 2H). In addition to the standard summary metrics generated by the Blackbox platform, we reanalyzed the tracking coordinate data to calculate novel parameters, including exploratory behavior (empty bin count), average speed, fore-to-hindpaw speed ratio, average acceleration, maximum acceleration, and acceleration standard deviation (Figure 2I-J, Supplemental Figure 2 associated with Figure 2). We found significant differences in empty bin counts, average speed and fore-to-hindpaw speed ratio (Figure 2I-J, Supplemental Figure 2D associated with Figure 2). Empty bin count, a metric representing the level of exploratory behavior and arena coverage, was consistently higher in SCI animals compared to sham across most time points (Figure 2I) (1 dpi *p* < 0.0001; 7 dpi *p* = 0.0002; 14 dpi *p* = 0.0009; 21 dpi *p* < 0.0001; 28 dpi *p* < 0.0001; 42 dpi *p* = 0.0021), indicating less exploratory behavior in SCI mice. Average speed (Figure 2J) was decreased across most time points in the SCI group compared to sham (1 dpi *p* = 0.0419; 7 dpi *p* = 0.0416; 21 dpi *p* = 0.0010; 28 dpi *p* = 0.0002; 42 dpi *p* = 0.0071). Such speed loss is reflective of the decreased motor activity exhibited by SCI mice and provides an important indicator of the extent of locomotor impairment in SCI animals. The fore-to-hindpaw speed ratio underscored the coordination deficits in SCI animals (Supplemental Figure 2D associated with Figure 2). At 1 dpi, the SCI group exhibited a significant decrease in this ratio compared to sham (*p* = 0.0071), indicating a loss of coordinated movement immediately following injury. The ratio remained relatively consistent in the SCI and sham groups after 1 dpi with a notable deviation at 35 dpi (*p* = 0.0328). No differences were found in average acceleration, maximum acceleration, or acceleration standard deviation (Supplemental Figure 2A-C associated with Figure 2). This indicates that the ability of mice to accelerate and the variability in that acceleration is not significantly affected by SCI under the conditions tested. These additional metrics provide deeper insight into the locomotor and behavioral differences between sham and SCI groups.

### Sustained Weight Distribution Alterations Following SCI

Utilizing luminance data, which reflects the pressure (or weight) placed onto each paw, we observed a persistent shift in weight distribution to the forepaws in SCI-treated animals. At baseline, both sham and SCI-treated animals exhibited uniform weight distribution across all limbs (Figure 3A). At 1 dpi, SCI-treated animals exhibited reduced hindlimb luminance intensity, indicative of decreased weight bearing on the hindlimbs. This reduction correlated with hindpaw dragging (Figure 3A, white arrows) and compensatory forelimb support (Figure 3A, white arrowheads). By 28 dpi, hindlimb weight-bearing improved in most animals, with weight placed on all four paws (Figure 3A). Analysis of the detected luminance-based features revealed that this post-injury shift in weight-bearing is primarily observed in fore and hindpaw luminances (Figure 3B). We analyzed the luminances in more detail to understand precisely how the animals altered their weight distribution after injury. While sham animals maintained stable weight distribution throughout the study, weight distribution still favored the forepaws after SCI as reflected in the average fore-to-hindpaw luminance ratio (Figure 3C). At 1 dpi, a significant shift to the weight-bearing on the forepaws is seen as an increased fore-to-hindpaw ratio in SCI-treated animals (*p* = 0.0041). Despite partial recovery, this fore-to-hindpaw ratio remained significantly elevated compared to sham controls until 42 dpi (*p* = 0.0007) (Figure 3C). The increase in the fore-to-hindpaw luminance ratio indicates a redistribution of pressure toward the forepaws, suggesting that SCI-treated animals shift their body weight forward.

**Figure 3.**
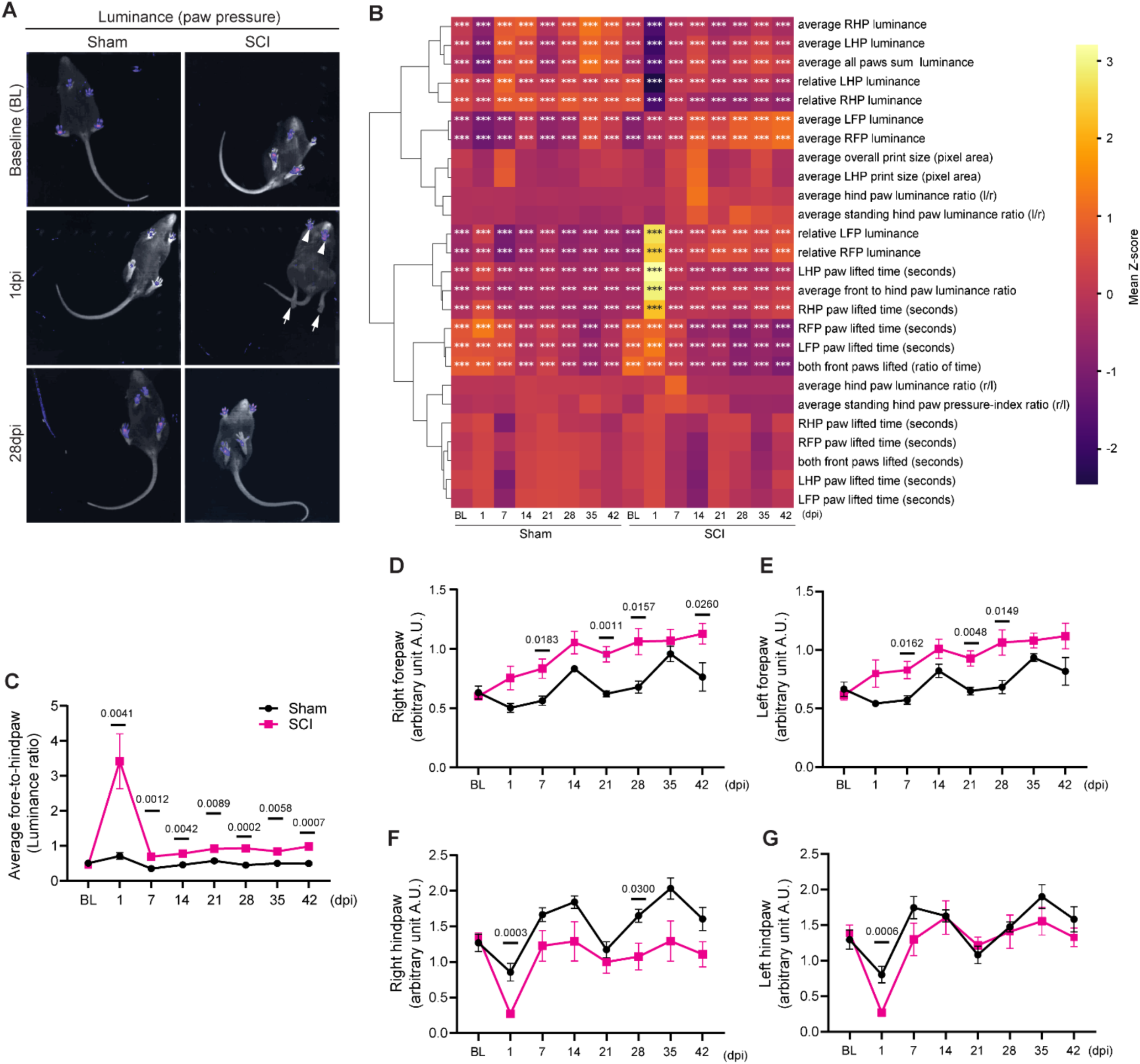
Changes in paw pressure following SCI. (A) Representative paw pressure luminance in Sham and SCI-treated animals at baseline (BL), 1-day post-injury (dpi), and 28 dpi, illustrating the distribution of weight-bearing across all four paws, with changes in intensity reflecting shifts in pressure distribution. (B) Hierarchical heatmap of the mean z-score for all detected features using luminance detection. Significance is represented as *** = *p*<0.001, ** = *p*<0.01, and * = *p*<0.05. Several luminance values were assessed during locomotion: (C) Average fore-to-hind paw luminance ratio over time. This ratio represents the relative pressure applied to the forepaws compared to the hindpaws. (D-E) Luminance of the right and left forepaws over time, measured as an indicator of weight-bearing on each forelimb. (F-G) Luminance of the right and left hindpaws over time, reflecting changes in hindlimb weight support. Data are shown as mean ± SEM for n = 6 Sham animals and n=8 SCI animals. Two outliers in the Sham group over three standard deviations from the mean were removed because of urine interfering with the luminance signal. Statistical significances using a two-way ANOVA are shown for (B) and using two-tailed unpaired Student’s t-test for (C-G) comparisons between Sham and SCI groups at specific time points.

Analysis of individual forepaw and hindpaw luminance data indicates that the increased fore-to-hindpaw ratio is largely driven by increases in forepaw pressure. The right and left forepaw luminances increased upon SCI (Figures 3D-E). These findings indicate increased weight-bearing on the forepaws following SCI. Conversely, the right and left hindpaw luminances significantly decreased at 1 dpi (*p* = 0.0003 and 0.0006, respectively), but were relatively unchanged for the remaining days post injury with the exception of the right hindpaw at 28 dpi (*p* = 0.0300) (Figures 3F-G). The sustained increase in pressure on the forepaws with relatively minimal decreased pressure on the hindpaws, highlights a persistent shift in weight distribution toward the forepaws immediately after SCI, which is maintained even after the animals have visually regained normal mobility.

### Principal Component Analysis of Behavioral Parameters Reflects Late-Stage Motor Recovery Differences between SCI and Sham Controls

Single-metric plots provide insight into specific parameter differences but lack a holistic representation of behavior. This limitation prevents a comprehensive understanding of overall changes in motor function. To address this, we implemented principal component analysis (PCA) to condense six selected significant motor metrics (i.e. fore and hindpaw distances, hindpaw angles (left and right), femur width, and distance traveled) into principal component space (Figure 4A). PCA components 1 and 2 capture the majority of the variance (Figure 4B). Additionally, to assess statistical significance between groups at each time point, we conducted multivariate analysis of variance (MANOVA).

**Figure 4.**
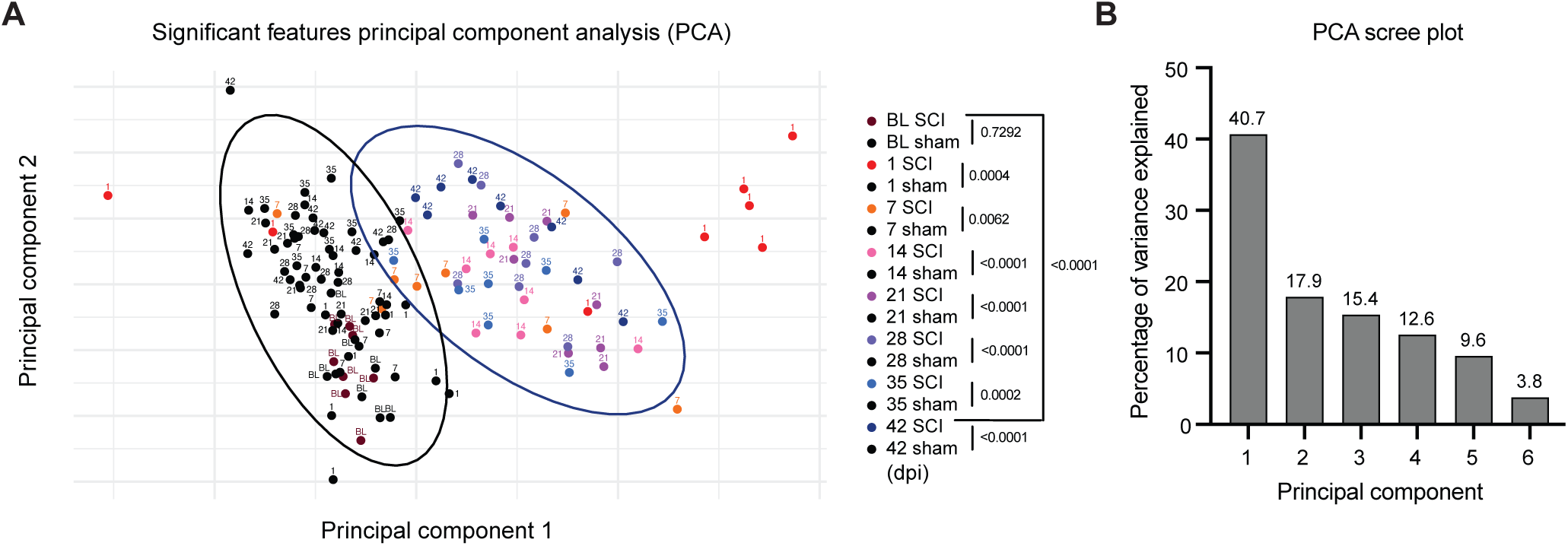
Principal Component Analysis (PCA) of locomotor and behavioral features following SCI. (A) PCA plot of selected significant features from individual mice across all time points. Each point represents a single mouse, with numbers and colors indicating the experimental group and time post-injury (see legend). The x-axis represents Principal Component 1 (PC1) and the y-axis represents Principal Component 2 (PC2), capturing the primary dimensions of variance in the dataset. Ellipses illustrate grouping trends among sham and SCI animals across different time points. (B) Scree plot showing the percentage of variance explained by the first six principal components. The x-axis represents the principal component number, and the y-axis represents the percentage of variance explained, indicating the relative contribution of each component to the overall variance in the dataset. Data are presented for n = 8 animals per group. Statistical significances using MANOVA are indicated between Sham and SCI groups across time points.

Throughout the study period, sham mice consistently clustered together across all time points, with baseline SCI mice localizing within a similar principal component (PC) space, suggesting uniform motor activity across uninjured animals. This observation was confirmed through MANOVA analysis, which showed no significant difference between the baseline sham and baseline SCI groups. At 1 dpi, SCI mice exhibited strong deviations in PC space localization, with many data points diverging substantially from both the sham and 7-42 dpi SCI clusters. This widespread distribution indicates a drastic and variable motor response to spinal cord injury. MANOVA analysis confirmed a statistically significant difference between SCI and sham animals at 1 dpi (*p* = 0.0004), further supporting this finding. By 7 dpi, although variability decreased, injured mice still exhibited dispersed localization within the PCA plot, reflecting persistent fluctuations in motor function and recovery rates (*p* = 0.0062). Differences between SCI and sham persisted for up to 42 dpi, with 7-42 dpi SCI forming distinct, separate cluster, indicating persistent motor differences despite visual motor recovery. These differences were validated through MANOVA, which demonstrated statistically significant difference between SCI and sham mice at each post-injury time point (14 dpi: *p* < 0.0001; 21 dpi: *p* < 0.0001; 28 dpi: *p* < 0.0001; 35 dpi: *p* = 0.0002; 42 dpi: *p* < 0.0001) (Figure 4A). Finally, MANOVA analysis confirmed a significant difference between SCI animals at baseline versus 42 dpi (*p* < 0.0001), indicating that even at the latest time point, SCI mice exhibited persistent motor impairment (Figure 4A). We also calculated effect sizes, which consistently revealed divergence between SCI and sham groups following injury (Baseline: 0.0474; 1 dpi: 0.6989; 7 dpi: 0.5424; 14 dpi: 0.7964; 21 dpi: 0.9656; 28 dpi: 0.8645; 35 dpi: 0.7315; 42 dpi: 0.8247). Therefore, by consolidating multiple metrics, PCA enabled us to uncover key recovery patterns that were not apparent when analyzing individual features in isolation. This approach underscores the importance of multi-metric data integration in capturing nuanced behavioral changes and advancing our understanding of spinal cord injury recovery.

### Keypoint MoSeq Facilitates Precise Identification and Analysis of Complex Motifs

We implemented Keypoint MoSeq (Weinreb et al., 2024) to identify nuanced, recurrent behaviors and extract deeper behavioral patterns from Blackbox-derived tracking data. This analytical approach provided an additional layer of behavioral detail, enabling the identification of subtle yet functionally significant changes in locomotion and motor patterns following SCI. MoSeq analysis enabled syllable-to-behavior mapping, through which we identified several distinct behavioral motifs (syllables) that were significantly less frequent among experimental groups (Control: sham & baseline SCI; 1 dpi SCI; 7-42 dpi SCI). These included forward acceleration, right and left turns and grooming behavior (Figure 5). Syllables 0, 1, 4, and 19 corresponded to forward acceleration trajectories, syllables 9, 10, and 11 to turn trajectories, and syllables 14 and 22 to grooming trajectories (Figure 5B). The following syllables were significantly decreased between Control and 1 dpi mice (syllable 0: *p* < 0.0001; syllable 1: *p* < 0.0001; syllable 4: *p* < 0.0001; syllable 10: *p* = 0.0145; syllable 11: *p* = 0.0284), and the following between the Control and 7-42 dpi groups (syllable 0: *p* < 0.0001; syllable 1: *p* < 0.0001; syllable 4: *p* < 0.0001; syllable 9: *p* = 0.0103; syllable 10: *p* = 0.0025; syllable 11: *p* < 0.0001; syllable 22: *p* = 0.0383). In addition, MoSeq can pick up on subtle behavioral differences between seemingly similar behavioral motifs. For example, syllables 14 and 22 are both designated as “grooming.” However, their syllable trajectories are slightly different (Figure 5B), resulting in syllable 22 being significantly different while syllable 14 is not. Similarly, syllables 0, 1, and 4 (forward acceleration) are significantly changed, while syllable 19 is not. The pattern across motifs indicates that animal motor activity is significantly impaired at 1 dpi, and despite apparent recovery, significant deficits persist into the later stages. By integrating behavioral motif frequency analyses, we uncovered distinct injury-related changes in movement execution.

**Figure 5.**
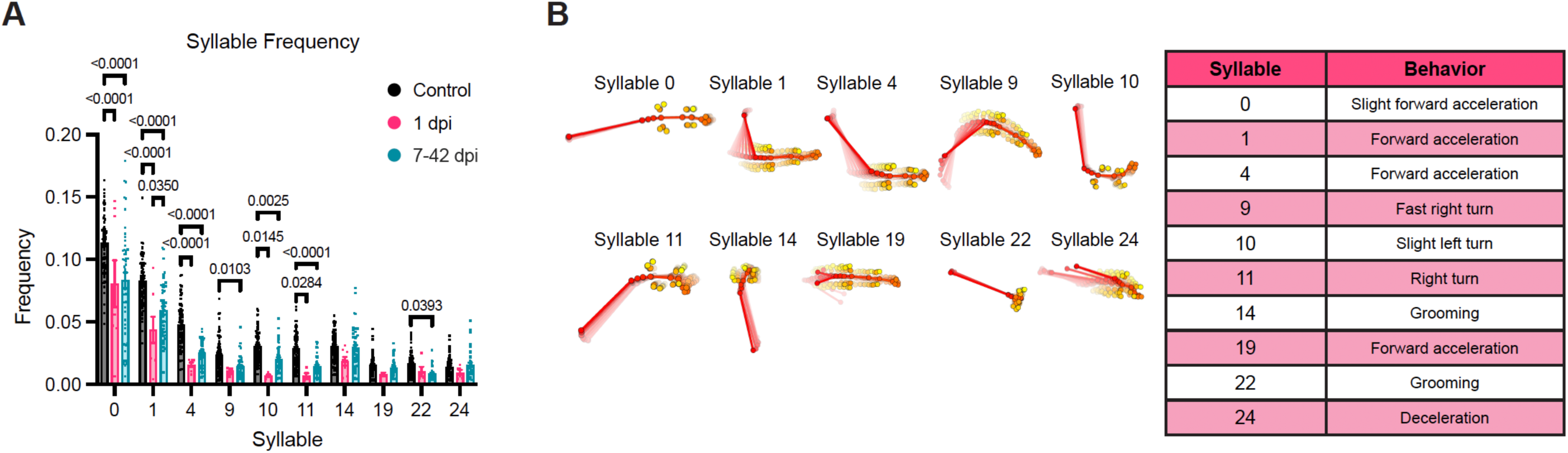
Keypoint MoSeq analysis reveals behavioral motifs and motor function patterns following SCI. (A) Frequency of selected behavioral motifs (syllables) in control, 1 dpi, and 7–42 dpi groups. The x-axis represents individual syllables corresponding to distinct movement patterns, while the y-axis represents the frequency of each syllable occurrence. (B) Syllable trajectory plots and syllable-to-behavior mapping. The left panel shows visualized movement trajectories associated with key syllables, depicting characteristic motion patterns. Different colored points represent different body parts, with orange indicating mouse head, red indicating central body parts and tail, and yellow indicating paws. The right panel presents a table mapping each syllable to its corresponding behavioral classification. Data are presented for n = 8 animals per group. Statistical significances using a two-way ANOVA with Tukey post hoc correction are shown for comparisons between Sham Control and SCI groups at specific time points.

### Blackbox Data Analysis Application

To provide users of Blackbox with an external application for more complex data analyses, we implemented the Shiny framework (Chang et al., 2025) using the R programming language (R Core Team, 2024) to develop a user-friendly, graphical user interface (Figure 6). Our data visualization and analysis tool provide a comprehensive suite of functionalities for the three types of Blackbox output data (Tracking, Features, and an aggregate summary .csv file). It enables the visualization of body-part coordinate-derived metrics such as speed plots (Figure 6A), feature-derived time series raster plots (Figure 6B), mouse trajectory visualizations plots (Figure 6C), and cage occupancy heatmaps (Figure 6D). Additionally, the tool facilitates simultaneous alignment and overlaying of these plots and allows users to download generated plots and data. Moreover, users can batch extract speed and feature metric averages, construct summary bar plots, and generate adaptable PCA plots (Figure 6E) for dimensionality reduction and visualization purposes.

**Figure 6.**
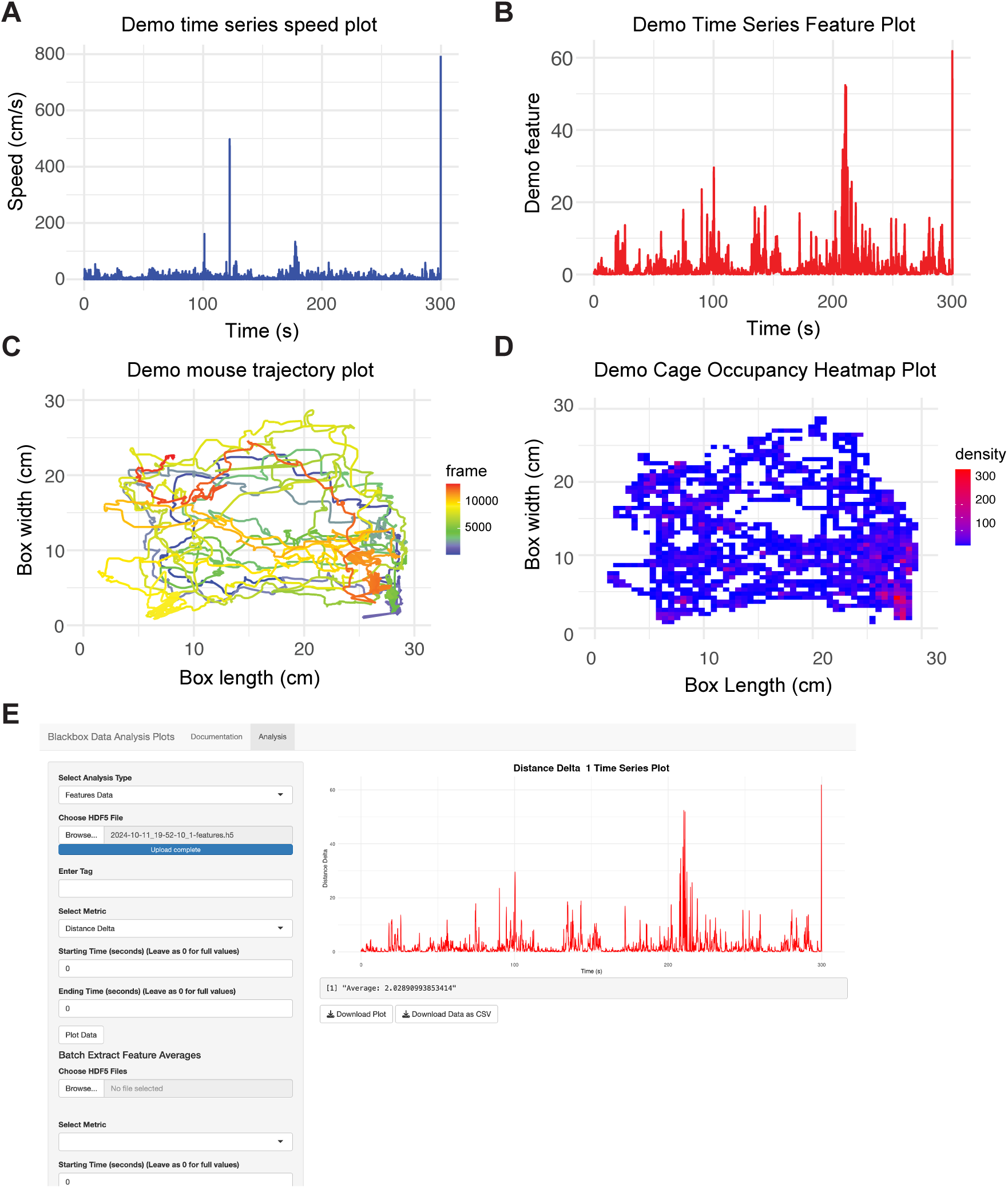
Blackbox Data Analysis App enables detailed, multifaceted behavioral data analysis. (A) Time series speed plot showing frame-by-frame analysis of animal speed over time. The x-axis represents time (s), and the y-axis represents speed (cm/s). (B) Time series feature plot displaying the progression of a selected behavioral feature across frames. The x-axis represents time (s), and the y-axis represents the feature value. (C) Mouse trajectory plot, visualizing movement paths within the testing environment. The x- and y-axes represent spatial dimensions (box length and width in cm), while color denotes the recording frame (blue = early in session, red = later in session). (D) Cage occupancy heatmap, showing spatial distribution of time spent in different areas of the enclosure. The x- and y-axes represent cage dimensions (cm), and the color scale represents occupancy density (blue = less time, red = more time). (E) Graphical user interface (GUI) of the Blackbox Data Analysis App, illustrating user input options for data selection, metric extraction, and visualization tools. This figure highlights the functionality of the BlackBox app for in-depth movement tracking and behavioral quantification.

By leveraging these comprehensive features, our tool enables researchers to seamlessly transform Blackbox system body-part tracking data and behavioral time-series data into intuitive visual representations without requiring prior coding experience. It allows for rapid assessment of mouse movement over the time course, serving as both a quality control measure and a way to identify novel features. This tool can enhance the interpretation of dynamic data patterns while facilitating deeper exploration of experiments.

### Thermal and Mechanical Sensitivity Returns to Baseline After SCI

To assess whether the postural adaptations detected via the Blackbox were due to pain, we tested thermal and mechanical sensitivity following SCI. Thermal sensitivity was evaluated using the Hargreaves assay at baseline and weekly from 7 to 42 days post-SCI. SCI-treated animals exhibited a significant increase in thermal withdrawal latency at 7 and 14 dpi in the average of left and right paw data, suggesting reduced thermal sensitivity, although motor impairment likely drives the increased withdrawal latency (*p* < 0.0001; Figure 7A). This increase in withdrawal latencies was observed bilaterally in the hindpaws, with the left paw having significantly higher latencies through 21 dpi and the right paw having a higher latency at 7 dpi (Figure 7B-C). The bilateral reduction in thermal withdrawal latency is consistent with the bilateral contusion model used (Figure 1). By 21 dpi, thermal withdrawal latency returned to baseline levels and remained stable through 42 dpi, indicating a full recovery of thermal sensitivity and ability to withdraw their hindpaw in SCI animals.

**Figure 7.**
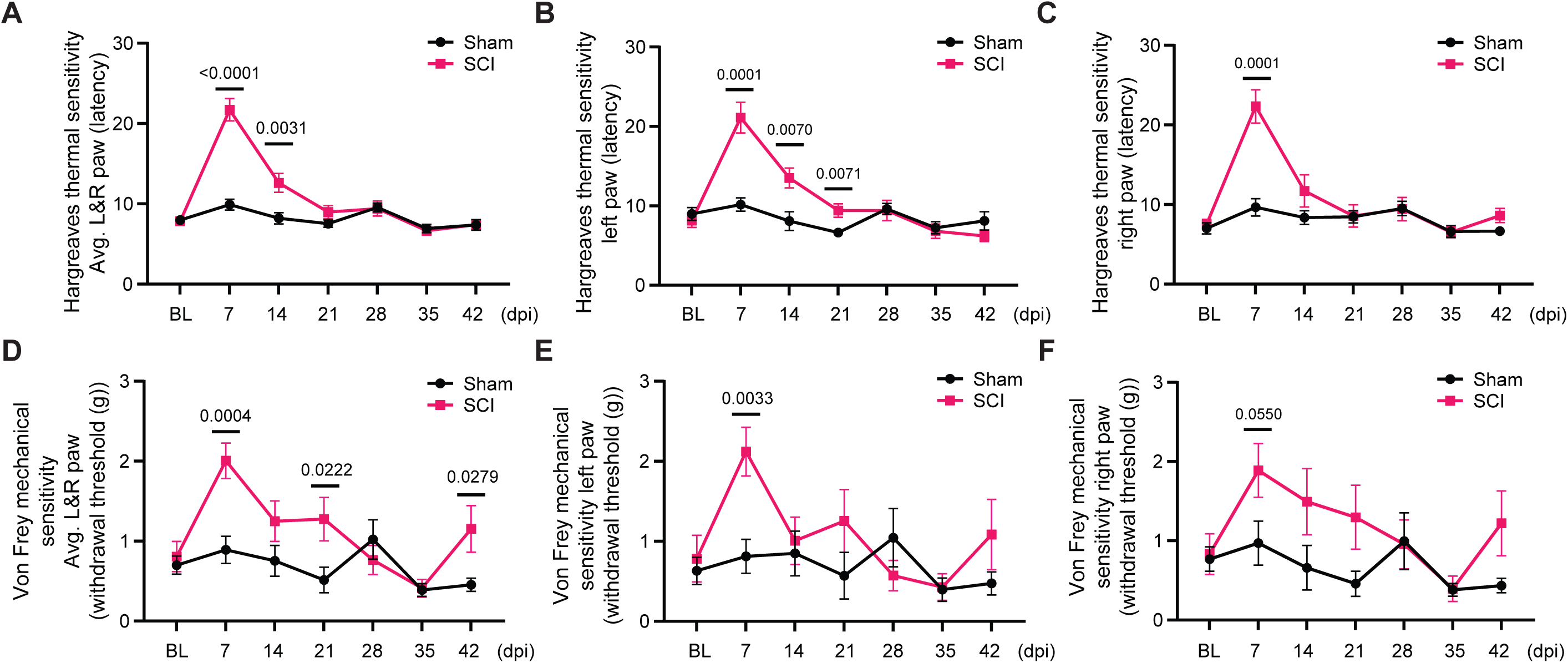
Recovery of Thermal and Mechanical Sensitivity Following SCI. (A, B, C) Thermal sensitivity recovery, measured by withdrawal latency (seconds) using the Hargreaves assay. (A) Average withdrawal latency, (B) left paw withdrawal latency, and (C) right paw withdrawal latency are shown over the recovery period. The x-axis represents days post-injury (dpi), and the y-axis represents latency in seconds. (D, E, F) Mechanical sensitivity recovery, measured by withdrawal thresholds (grams) using the simplified up-down (SUDO) Von Frey method. (D) Average withdrawal threshold, (E) left paw withdrawal threshold, and (F) right paw withdrawal threshold are shown over the recovery period. The x-axis represents days post-injury (dpi), and the y-axis represents withdrawal threshold in grams. Data represent n = 8 animals per group, with mean ± SEM displayed. Statistical significances using two-tailed unpaired Student’s t-tests are shown for comparisons between Sham and SCI groups at specific time points.

Mechanical sensitivity was assessed using the Simplified Up-Down (SUDO) Von Frey method at the same timepoints (Bonin et al., 2014). Similar to thermal sensitivity, SCI-treated animals displayed a significant increase in withdrawal threshold to mechanical stimuli (Figure 7D), suggesting reduced mechanical sensitivity. However, similar to the thermal sensitivity assay, this increased withdrawal threshold is likely driven by motor impairment. At 7 dpi, the average withdrawal threshold of both hindpaws is significantly increased in SCI mice (*p* = 0.0004, Figure 7D), which can be seen individually in both the left (*p* = 0.0033, Figure 7E) and right hindpaws (*p* = 0.0550, Figure 7F). In contrast to thermal withdrawal latencies, mechanical withdrawal thresholds were variable over the time course of recovery.

## Discussion

By integrating high-resolution kinematic analysis, machine learning-based pose estimation, and weight distribution metrics, our study identified persistent locomotor impairments and compensatory adaptations that were not detected by traditional behavioral assays. Notably, SCI-induced alterations, such as widened forepaw and reduced hindlimb spacing, altered hindpaw angles, increased forelimb weight-bearing, and diminished locomotor activity, persisted beyond the typical recovery period, indicating long-term neuromuscular and biomechanical adjustments. This demonstrates the importance of utilizing highly quantitative and machine-learning approaches for evaluating motor and postural recovery following SCI.

### Advantage of Quantitative Motor Behavior Assessments

Our findings build upon previous research utilizing conventional SCI assessment tools such as the BMS locomotor scale and CatWalk system. While these approaches provide valuable insights into gross motor function, they often lack the sensitivity to capture nuanced movement deficits (Basso et al., 2006; Hamers et al., 2006). The BMS scale is limited by inter-rater variability and an inability to detect subtle locomotor compensations (Ito et al., 2022), whereas CatWalk is limited by its constrained pathway, which may not reflect natural movement dynamics (Walter et al., 2020). By employing high-speed video tracking, machine learning-based body part estimation, and weight-bearing metrics, we overcame these limitations and identified sustained postural adaptations and gait abnormalities that persisted beyond the expected recovery period. Our findings are consistent with recent studies showing that deep learning-based kinematic analysis can reveal subtle locomotor deficits overlooked by traditional assessments such as the BMS (Ascona et al., 2024; Eisdorfer et al., 2025). Ascona et al. found that the average mouse speed decreased after SCI, similar to our findings, and Eisdorfer et al. used MoSeq to identify changes in behavioral motifs during the SCI recovery period. Indeed, quantitative motor assessments after SCI will soon become the gold standard. Our study complements these studies by leveraging body-part tracking data to quantitatively capture logical postural and weight-bearing metrics that change post-SCI.

### Sensory Recovery Post-SCI

Human patients can experience a loss of sensation and the development of allodynia and hyperalgesia after SCI (Finnerup et al., 2001; Siddall & Loeser, 2001). While thermal and tactile allodynia has been reported in rats post-SCI (Detloff et al., 2013; Lindsey et al., 2000), studies in mice yield conflicting results, with some strains exhibiting allodynia and others not (Detloff et al., 2012). Mouse strain-dependent variability and injury severity may explain some of the disagreement in sensory recovery. BALB/c mice develop tactile allodynia and spontaneous below-level pain behaviors post-SCI (Kerr & David, 2007). C57BL/6 mice are also reported to develop thermal and tactile allodynia; however, one study found tactile allodynia developed only after severe contusion injury (Hoschouer et al., 2010; Kerr & David, 2007; Lee et al., 2023; Yang et al., 2024).

We find that thermal and mechanical sensitivity recovers to baseline without subsequent hypersensitivity in C57BL/6J female mice. Even though we do not detect hypersensitivity after SCI, our results align with previous findings that thermal sensitivity typically recovers more reliably than mechanical sensitivity, which exhibits greater variability (Hoschouer et al., 2010). This difference could arise from distinct neural mechanisms, particularly within the spinal cord, that differentially influence the recovery of thermal and mechanical sensitivity following SCI (Berkey et al., 2020; Gwak et al., 2017). Notably, studies have shown that persistent mechanical hypersensitivity following SCI may result from maladaptive plasticity within dorsal horn circuits, including disinhibition and reorganization of excitatory and inhibitory neuronal networks (Garraway & Huie, 2016).

These findings underscore the need for comprehensive, multimodal sensory assessments to fully capture post-injury sensory dysfunction and highlight the limitations of evoked pain behaviors as reliable indicators of neuropathic pain in mouse SCI models. Furthermore, studies using mouse grimace scales (MGS) have suggested transient pain-like behaviors primarily within the first week post-injury, with minimal chronic hypersensitivity (Heinsinger et al., 2020; Langford et al., 2010). These uncertainties evaluating pain perception in mice reinforce the necessity of refining behavioral pain assessments and integrating additional physiological measures, such as calcium imaging and electrophysiological recordings, to provide a more comprehensive evaluation of pain processing after SCI.

### Behavioral Motif Analysis

Beyond traditional motor and sensory analyses, we leveraged advanced computational techniques, including Keypoint MoSeq (Weinreb et al., 2024) and PCA-based population analyses, to extract deeper behavioral insights. MoSeq revealed distinct shifts in locomotor motifs across injury groups, such as decreased frequency of forward accelerations and turns, highlighting persistent post-injury changes. This aligns with previous studies demonstrating that rodents with SCI modify their movement patterns to maintain stability and mitigate functional loss (Hansen et al., 2012). Similarly, a recent study using various SCI models also found behavioral motif differences in functional recovery using MoSeq (Eisdorfer et al., 2025). Interestingly, in our study, even though average acceleration, maximum acceleration, and acceleration standard deviation were unchanged, MoSeq identified that the forward acceleration motifs were less frequent. Again, highlighting the benefit of machine-learning algorithms to identify subtle behavioral changes. Meanwhile, PCA revealed that despite improvements in gross locomotion, SCI-treated animals retained distinct motor profiles separate from Sham controls, underscoring persistent postural and neuromuscular differences (Krizsan-Agbas et al., 2014). This is consistent with prior reports showing that, although behavioral improvements are observed following SCI, underlying motor control deficits often persist, indicating incomplete functional recovery.

### Study considerations

While our study provides a comprehensive evaluation of SCI recovery, several considerations should be acknowledged. First, our analysis was conducted in a single mouse strain and injury severity, which may limit generalizability to other models. Future studies should explore whether these findings hold across different genetic backgrounds and injury paradigms (Sharif-Alhoseini et al., 2017). Investigating whether similar postural adaptations and compensatory mechanisms occur across diverse injury paradigms, including different severities of contusion, hemisection, and transection SCI models, will enhance the generalizability of our findings (Cheriyan et al., 2014). Additionally, expanding analyses to different mouse strains and sexes will provide insight into potential genetic and sex-specific differences in recovery (Farooque et al., 2006; Nishi et al., 2020; Sipski et al., 2004). While the Blackbox platform excels in detecting postural and gait abnormalities, integrating complementary techniques such as electromyography (EMG) could further elucidate neuromuscular recovery dynamics (Hansen et al., 2012). Lastly, long-term follow-up beyond 42 dpi would provide insight into whether these postural adaptations eventually normalize or persist indefinitely (Battistuzzo et al., 2016).

## Conclusion

Our results provide valuable translational insight into the subtle parameters changed in mouse SCI recovery. Careful quantitative analyses of SCI recovery can objectively detect subtle motor deficits, providing a valuable tool for evaluating the efficacy of experimental therapies like neuroprotective drugs, biomaterials, and stem-cell-based therapies (Fawcett et al., 2007).

## Acknowledgements

We thank Kyle Baumbauer for insights into spinal cord injury and Seungwon (Sebastian) Choi for introduction to the Blackbox system. This work was supported by NIH grants R01NS127575, R01NS117065, and R01NS111776 to C.L.Z. and R21NS128203 and R01MH134799 to H.C.L.

## Materials and Methods

### Animals

Female wild-type C57BL/6J mice (JAX: 000664; RRID: IMSR_JAX:000664), aged two months old at time zero of the experiment, were used in all experiments. Female mice were used because they are anatomically easier to manage for post-operative bladder expression, which is critical for survival and consistent care after SCI, whereas male mice are more prone to urinary obstruction and require more intensive care. Mice were housed in a temperature-controlled facility under a 12-hour light/dark cycle with unrestricted access to food and water. Sample sizes were determined empirically, and all animal procedures were approved by the Institutional Animal Care and Use Committee (IACUC) at UT Southwestern Medical Center.

### Spinal Cord Injury Model

Adult mice were anesthetized using a ketamine (100 mg/kg) and xylazine (10 mg/kg) cocktail, followed by a laminectomy at the thoracic level 9 (T9) spinal segments to expose the spinal cord. Mice were assigned to either the Sham or SCI group, with each group consisting of eight animals. For the SCI group, a standardized contusion injury was induced using the IH impactor (Precision Systems and Instrumentation, Lexington, KY; Model IH-0400) with a 1-mm-diameter impactor tip, applying a force of 60 kdyn perpendicular to the central axis of the exposed dorsal spinal cord. The tip was immediately retracted after impact to prevent excessive tissue damage. Sham animals underwent identical surgical procedures, including laminectomy and impactor tip positioning, but no injury was delivered (Tai et al., 2021). This contusion model produces a bilateral lesion with a central cavity surrounded by spared tissue, mimicking clinically relevant traumatic spinal cord injuries. Postoperatively, all mice were monitored in their home cages and provided supportive care, including manual bladder expression twice daily until reflexive bladder control was regained.

### Tissue Processing and Immunohistochemistry

Mice were euthanized via CO₂ overdose and transcardially perfused with ice-cold phosphate-buffered saline (PBS), followed by 4% paraformaldehyde (PFA) in PBS. The entire spinal cord was carefully extracted, post-fixed in 4% PFA overnight at 4°C, and cryoprotected in 30% sucrose at 4°C. A 1.5-cm segment encompassing the injury site was sectioned at 20 µm using a Leica cryostat. Immunohistochemistry was performed following a previously established protocol (Wang et al., 2016). Tissue sections were incubated overnight at 4°C with primary antibodies, including chicken anti-GFAP (1:1000, Abcam, ab4674) and rabbit anti-PDGFR beta (1:500, Abcam, ab32570), diluted in blocking buffer containing 3% goat serum in 0.3% PBST. Sections were then incubated with secondary antibodies for two hours at room temperature, including donkey anti-chicken Alexa Fluor 488 (1:1000, Thermo Fisher Scientific, A78948) and donkey anti-rabbit Alexa Fluor 555 (1:1000, Thermo Fisher Scientific, A31572). Confocal imaging was performed using a Nikon A1R confocal microscope with a 20× objective, and image stitching was carried out using NIS-Elements software. ImageJ was used for fluorescence area analysis.

### Behavioral Assessments

All behavioral assessments were conducted by a single experimenter blinded to group assignments. To ensure consistency, animals were acclimated to the testing environment in two sessions: a one-hour habituation period on the day before testing and an additional 30-minute session immediately prior to testing. Sensory assays, including mechanical and thermal sensitivity tests, were performed at baseline (7 days prior to SCI) and weekly from 7 to 42 days post-injury (dpi). Motor assessments were conducted at baseline (7 days prior to SCI), 1 dpi, and then weekly until 42 dpi.

### Von Frey Mechanical Sensitivity Testing

Mechanical sensitivity was evaluated using the Simplified Up-Down (SUDO) method (Bonin et al., 2014) to determine withdrawal thresholds to calibrated Von Frey filaments. Mice were acclimated to an elevated plastic chamber with a mesh floor, allowing access to the plantar surface of the hind paws. Filaments ranging from 0.02 g to 1.4 g were applied to the left and right hind paws, and a positive response was recorded if the animal exhibited behaviors such as toe spreading, flinching, licking, or paw withdrawal. To minimize sensitization, a five-minute interval was maintained between applications.

### Hargreaves Thermal Sensitivity

Thermal sensitivity was evaluated using the Hargreaves method, where animals were acclimated to a plastic chamber positioned on a heated (30°C) glass platform before testing (Hargreaves et al., 1988; Landy et al., 2021). A focused beam of light was directed at the plantar surface of the hind paws to measure withdrawal latency, with beam intensity calibrated to elicit a response within approximately 10 seconds in uninjured wild-type animals. Each hindpaw was tested three times with a 5-minute interval between tests to ensure consistency. A cutoff time of 30 seconds was implemented to prevent potential tissue damage in cases of delayed withdrawal.

### Motor Recovery Analysis

Motor behavior and recovery were assessed using the BlackBox One motion tracking system (Zhang et al., 2022) in combination with Palmreader v0.7.0 and Analysis v0.3.0 software. Animals were placed in a 6 × 6-inch opaque enclosure and recorded from below for 5 minutes under low-light conditions. The system utilized frustrated total internal reflection (FTIR) technology and Optitouch sensors to detect luminance in response to surface contact, which corresponded to the pressure applied by each limb. Concurrently, video-based pose estimation was used to track body position and movement. Palmreader software generated summary files containing averaged metrics such as paw luminance, limb separation, and limb angles, while detailed tracking data was stored in h5 format for further analysis. To encourage continuous movement, animals were not acclimated to the testing chamber before recording.

Significant features for body tracking and luminance data were analyzed using Python code provided by BlackBox One (Cole et al., 2025). The z-score for each feature was calculated using the standard deviation and mean values from both Sham and SCI across all timepoints. Significance was determined using a two-way ANOVA and Bonferroni adjusted p-values are reported.

### Skeleton Mislabeling Analysis

To evaluate the accuracy of skeletal pose estimation in the BlackBox system following SCI, a manual frame-by-frame analysis of skeleton data was conducted. Videos generated by Palmreader were converted into individual JPEG frames at a rate of one frame per second. Mislabeled skeleton components were identified and quantified based on predefined criteria, including (1) incorrect connections between body parts, such as a forepaw linked to a hind paw; (2) lines appearing in empty space without corresponding body parts, such as a misidentified tail position; and (3) exaggerated skeletal extensions, such as toe lines extending beyond the anatomical boundaries of the limb.

### Tracking Features Generation

To develop novel movement metrics from the BlackBox tracking dataset, we implemented custom scripts in the R programming language. The tailbase coordinate was used as the primary reference point for all tracking analyses due to its high reliability. The complete code for these analyses is available at GitHub (https://github.com/helenLaiLab/BlackBox_Analysis/tree/main/tracking_metric_generation.).

(1) Speed Metrics — The Euclidean distance formula (*Distance* = √[(x₂ – x₁)² + (y₂ – y₁)²]) was used to calculate the displacement of the tailbase coordinate between consecutive frames, which was then divided by the frame interval to determine frame-by-frame speed. The mean speed was computed by averaging all individual frame speeds. Additionally, the fore-to-hind paw speed ratio was determined by averaging the speeds of both forepaws and both hind paws separately and calculating the ratio of forepaw speed to hindpaw speed.
(2) Acceleration Metrics — Acceleration was calculated by determining the rate of change in speed over time, yielding frame-by-frame acceleration values. From these data, the average acceleration, maximum acceleration, and standard deviation (acceleration variability) were computed.
(3) Empty Bins Count — The recording area of the BlackBox enclosure was divided into 2,500 equally sized bins, each with an area of 0.36 cm². The bin number for each recorded coordinate was determined, and the total number of bins without any recorded coordinates was summed to obtain the empty bins count.

### Principal Component Analysis (PCA)

PCA was conducted using the R programming language to analyze selected significant features derived from Palmreader software and additional tracking-based metrics. The analyzed features included fore and hindpaw distances, hindpaw angles (left and right), femur width, and distance traveled. To generate the PCA plot, the selected features were averaged across all mice and time points, and the `prcomp()` function in R was used to compute the principal components. Principal components 1 and 2 were plotted on the x- and y-axes, respectively, with ellipses indicating time-point clusters. To assess statistical differences between clusters in PCA space, a multivariate analysis of variance (MANOVA) was performed using the `manova()` function in R, comparing sham and SCI groups at each time point. The resulting p-values were visualized in a bar plot. Effect size was calculated using Pillai’s Trace, a statistic included in the MANOVA analysis. The complete PCA generation script is available at GitHub (https://github.com/helenLaiLab/BlackBox_Analysis/tree/main/PCA).

### BlackBox Data Analysis App

A user-friendly data analysis application was developed using the Shiny framework in R to facilitate processing and visualization of BlackBox motion tracking data. The app is designed to handle various file types outputted by BlackBox and enables detailed behavioral and locomotor analyses. The raw scripts are available at GitHub (https://github.com/helenLaiLab/BlackBox_Analysis/tree/main/BlackBox_Plotting_Raw_Scripts).

The application converts the tracking HDF5 file into an R data frame using the rhdf5 package (Fischer et al., 2024), allowing extraction of key locomotor features, including:

(1) Speed — Calculated as described in the speed metrics section.
(2) Likelihood — Extracted values representing the confidence of coordinate estimates from the tracking data.
(3) Trajectory — Visualization of movement paths, color-coded by time to reflect changes over the recording period within a scaled 15.24 x 15.24 cm arena.
(4) Cage Occupancy Heatmap — Generated by binning coordinate values into 2,500 regions covering the recording area, with color intensity indicating time spent in each region. Transparent areas denote regions where no movement was recorded.
(5) Batch Extraction of Speed Averages — Computes and exports speed metrics for selected body parts, summarizing results into an Excel file.

The app also processes BlackBox feature HDF5 files, extracting and renaming feature metrics for improved readability. It generates:

(1) Feature Plots — Time-series line or raster plots visualizing numeric and ethogramic (Boolean) data.
(2) Batch Extraction of Feature Averages — Exports averaged feature values per file into an Excel summary.

Additionally, the app supports importing BlackBox summary CSV files for generating bar plots and allows PCA summary file imports for on-demand PCA plot generation. Data preprocessing is performed with the dplyr (Wickham et al., 2023), readxl (Wickham & Bryan, 2023), jsonlite (Ooms, 2014), writexl (Ooms, 2024), and stringr (Wickham, 2023) packages. Data visualization is performed using the colourpicker (Attali, 2023), ggplot2 (Wickham, 2016), patchwork (Pedersen, 2024), randomcoloR (Ammar, 2019), rgl (Murdoch & Adler, 2024), shinyjs (Attali, 2021), shinyWidgets (Perrier et al., 2024) packages.

### Motion Sequencing Analysis (MoSeq)

To identify nuanced behavioral motifs, the Keypoint Motion Sequencing (MoSeq) platform was utilized. Analysis was performed using the MoSeq team’s Jupyter Notebook via Google Colab. The customized Colab notebooks and configuration files are available at GitHub (https://github.com/helenLaiLab/BlackBox_Analysis/tree/main/MoSeq).

Input data included tracking HDF5 files and tracking-AVI videos generated by BlackBox. The dataset was configured following the “Option 1: Setup from DeepLabCut” approach, and a subset of 19 tracking files was randomly selected for model training. PCA was performed, followed by hyperparameter tuning to determine an optimal kappa value of 10^10^. The dataset was then modeled using an Autoregressive Hidden Markov Model (AR-HMM), and behavioral syllables were sorted by frequency before exporting model outputs to CSV. Given the dataset size, the trained model was applied in a batch-wise manner, splitting the dataset into two halves for analysis before merging results. MoSeq-generated visualizations included trajectory plots, grid movies, and syllable similarity dendrograms. For statistical analysis, data were categorized into three groups: (1) control (sham and baseline SCI), (2) 1 dpi SCI, and (3)7-42 dpi SCI, to improve interpretability while maintaining behavioral relevance. The statistical analysis followed the MoSeq Jupyter Notebook pipeline, generating plots and summary data frames. For syllable frequency analysis, the MoSeq stats_df data frame was used to identify the ten syllables with the highest overall variability across all metrics. Variance was determined using weighted standard deviation calculations in R, with values summed across metrics for each syllable. MoSeq-generated trajectory plots were examined to assign behavioral classifications to syllables, informed by corresponding grid movies.

### Statistical Analysis

All statistical analyses were conducted using R and GraphPad Prism v10.4.1. Comparisons between Sham and SCI groups were performed using unpaired, two-tailed Student’s t-tests for single-variable comparisons. For multi-variable and time-series data, two-way ANOVA with Tukey post hoc corrections was applied to control for multiple comparisons. Data are presented as mean ± standard error of the mean (SEM). Statistical significance was defined as *p*<0.05, and figures were generated using GraphPad Prism to visualize recovery trajectories and group differences.

---

**Supplementary Figure 1.**
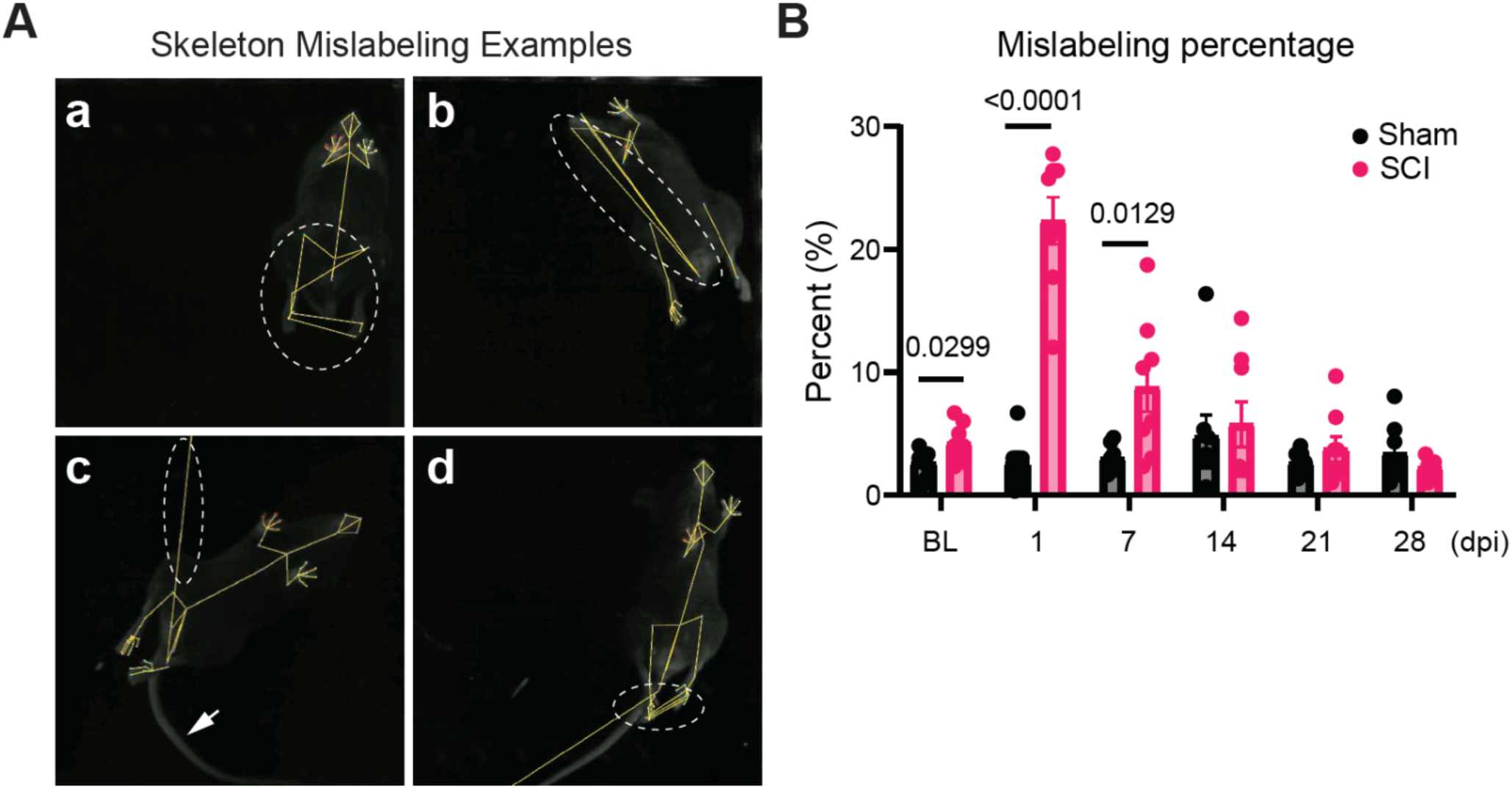
associated with Figure 2. Skeleton mislabeling in SCI-treated animals. (A) Examples of skeleton mislabeling observed in SCI-treated animals. (a, b) Incorrect connections between body parts (dashed circles, video 1 seconds 5 & 80). (c) Misidentification of the tail, with the actual tail position indicated by a white arrow (video 2 second 65). (d) Extensions beyond body boundaries, such as exaggerated toe lengths (dashed circle, video 3 second 5). (B) Quantification of mislabeling frequency in Sham and SCI-treated animals over time. The x-axis represents days post-injury (dpi), while the y-axis represents the percentage of mislabeling occurrences. Data are presented as mean ± SEM for n = 8 animals per group. Statistical significances using two-tailed unpaired Student’s t-tests are shown for comparisons between Sham and SCI groups at specific time points.

**Supplementary Figure 2.**
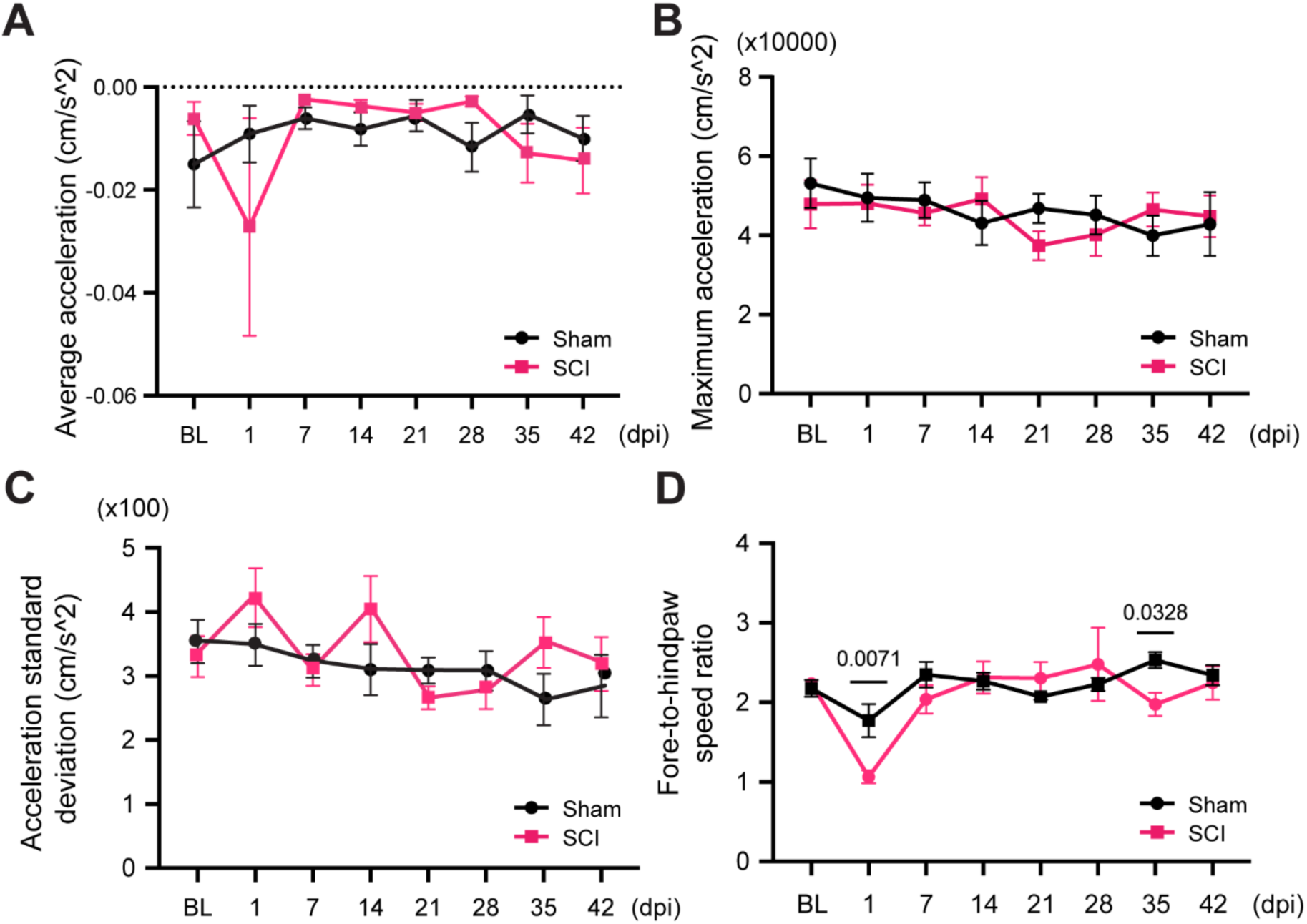
associated with Figure 2. Novel metrics assessing acceleration following SCI. (A) Average acceleration (cm/s²) over time. (B) Maximum acceleration (cm/s²) over time, representing peak recorded acceleration during the observation period. (C) Acceleration variability (standard deviation, cm/s²), representing fluctuations in movement speed over time. (D) Fore-to-hind paw speed ratio, an indicator of interlimb coordination during gait. Data are presented as mean ± SEM for n = 8 animals per group. Statistical significances using a two-way ANOVA with Tukey post hoc correction are shown for comparisons between Sham and SCI groups at specific time points.

